# CRAFT: Compact genome Representation towards large-scale Alignment-Free daTabase

**DOI:** 10.1101/2020.07.10.196741

**Authors:** Yang Young Lu, Jiaxing Bai, Yiwen Wang, Ying Wang, Fengzhu Sun

**Author notes:** Co-first authors (Yang Young Lu, Jiaxing Bai).

## Abstract

**Motivation:** Rapid developments in sequencing technologies have boosted generating high volumes of sequence data. To archive and analyze those data, one primary step is sequence comparison. Alignment-free sequence comparison based on *k*-mer frequencies offers a computationally efficient solution, yet in practice, the *k*-mer frequency vectors for large *k* of practical interest lead to excessive memory and storage consumption.

**Results:** We report CRAFT, a general genomic/metagenomic search engine to learn compact representations of sequences and perform fast comparison between DNA sequences. Specifically, given genome or high throughput sequencing (HTS) data as input, CRAFT maps the data into a much smaller embedding space and locates the best matching genome in the archived massive sequence repositories. With 10^2^ – 10^4^-fold reduction of storage space, CRAFT performs fast query for gigabytes of data within seconds or minutes, achieving comparable performance as six state-of-the-art alignment-free measures.

**Availability:** CRAFT offers a user-friendly graphical user interface with one-click installation on Windows and Linux operating systems, freely available at https://github.com/jiaxingbai/CRAFT.

**Supplementary information:** Supplementary data are available at *Bioinformatics* online.

## 1 Introduction

Rapid developments in efficient and affordable sequencing technologies have enabled biologists to collect unprecedented high volumes of sequence data. For example, the size of NCBI RefSeq Database (Pruitt *et al.*, 2005) reaches the magnitude of terabyte (TB). Considering the massive collection of genomic sequences, one fundamental problem related to many biological studies is sequence comparison within the collection (Perelman *et al.*, 2011; Wood and Salzberg, 2014; Lu *et al.*, 2017b), along with the technical challenges about how to organize and transmit the data promptly. The workhorse for sequence comparison is sequence alignment (Altschul *et al.*, 1990), where global alignment emphasizes the entirety (Needleman and Wunsch, 1970) and local alignment considers only local regions of high similarity (Smith *et al.*, 1981). However, these alignment-based methods are not only computationally expensive, but also relying on large-size indexing that is not scalable to massive sequence repositories.

Alignment-free methods (Lu *et al.*, 2017a; Zielezinski *et al.*, 2017, 2019) are other alternatives that are much more computationally efficient than conventional alignment-based methods, and have been widely applied to genomic and metagenomic comparisons (Bernard *et al.*, 2018; Wang *et al.*, 2014). Most alignment-free methods compare the dissimilarity of sequences in terms of the *k*-mer frequency vectors, where *k*-mers refer to the set of words tokenized from the sequence of fixed length *k*. However, the popularity of those methods, including 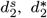 (Lu *et al.*, 2017a), CVTree (Qi *et al.*, 2004), co-phylog (Yi and Jin, 2013), etc., is limited because the *k*-mer frequency vectors for large *k* of practical interest result in excessive memory and storage consumption. One alternative to tackle the scalability of alignment-free methods is by hashing, such as Mash (Ondov *et al.*, 2016) and Skmer (Sarmashghi *et al.*, 2019), which use MinHash to approximate Jaccard distance between *k*-mer presence/absence vectors of pairwise sequences. However, the presence/absence of *k*-mers is less informative compared to using *k*-mer frequencies (Lu *et al.*, 2017a; Ren *et al.*, 2018).

In this paper, we propose CRAFT, a general tool for compact representations of gigantic sequence databases and fast genomic/metagenomic sequence comparison. Based on the co-occurrences of adjacent *k*-mer pairs, CRAFT maps the input sequences into a much smaller embedding space, where CRAFT offers fast comparison between the input and pre-built repositories. Notably CRAFT archives three widely-used genomic/metagenomic sequence repositories, including 139, 576 genomes from NCBI Assembly, 7,106 representative genomes from RefSeq, and 2, 355 meta-genomic samples from Human Microbiome Project (HMP 1-II), reducing the file size from 829,60GB (.fa), 376.38G (.fa), and 7.13TB (.sra) to 2.93GB, 152.67MB, and 50.40MB, respectively. From practitioners’ perspective, CRAFT offers a userfriendly graphical user interface with one-click installation on Windows and Linux operating systems. We evaluate the performance of CRAFT from multiple perspectives, in comparison to state-of-the art alignment-free measures including Manhattan, CVTree, 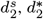, Mash, and Skmer. (1) CRAFT is used to estimate pairwise dissimilarities for datasets of different diversities, including 28 vertebrate assembled genomes, 27 *E. coli* and *Shigella* assembled genomes, 21 primate genomes and 28 metagenome samples from mammal guts. Evaluated by consistency to evolutionary distance or reference grouping tree, CRAFT demonstrates its superiority particularly in datasets with high diversity (2) CRAFT is used to search two high throughput sequencing (HTS) datasets against archived massive sequence repositories. In the first dataset, 104 randomly selected HTS data from 8 major taxonomic branches are queried against NCBI assembled genomes, and 90 out of 104 can locate their source genomes within top-2 matches. In the second dataset, 13 randomly selected HTS metagenomic are queried against HMP 1-II, and 11 out of the 13 can be matched to their targeted body locations exactly (Section 3.3). (3) The robustness of CRAFT is demonstrated in the sense that even incomplete genome sequences (e.g., 5% of sequence) and low-depth shotgun sequencing data (e.g., 1% coverage of reads) can obtain accurate searching performance (Section 3.4).

## 2 Methods

### 2.1 Workflows

As shown in Figure 1, CRAFT includes two key processing steps: *embedding* and *dissimilarity.* Given a genomic/metagenomic sequence as input, *embedding* learns its compact representation by mapping nucleotide sequences into low-dimensional space. In brief, CRAFT counts the co-occurred *k*-mers of the query sequence with 1–bp step-size sliding windows. Based upon the co-occurrence of adjacent *k*-mers, each *k*-mer is transformed to a vector using the GloVe algorithm (Pennington *et al.*, 2014). After that, *dissimilarity* calculates the dissimilarities between the query sequence and the sequences in the archived repositories, in terms of their learned compact representations. Additionally, CRAFT provides three types of built-in downstream visualized analyses of the query results, including (1) clustering the sequences into dendrograms using the UPGMA algorithm (Murtagh, 1984); (2) qualitatively eyeballing the dissimilarities of sequences by the bars; (3) embedding the dissimilarities of sequences spatially in two-dimensional spaces by principal coordinate analysis (PCoA) (Gower, 1966).

**Fig. 1.**
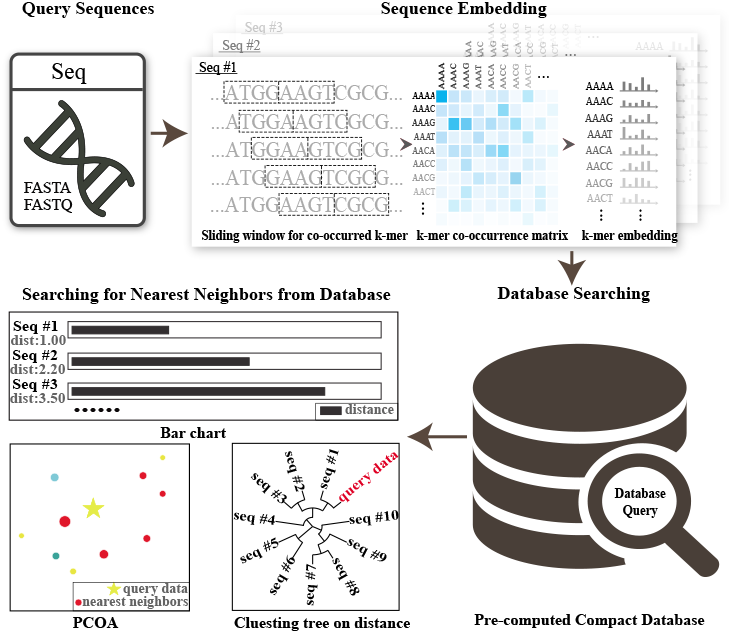
The workflow of CRAFT. CRAFT parses the input genomic/metagenomic sequences, counts the co-occurrences of adjacent *k*-mer pairs, learns the embeddings, searches against archived repositories by the dissimilarity between the input and the archived sequences in the embedding space, and finally reports the best matches in various ways.

### 2.2 CRAFT *embedding*

CRAFT takes genomic/metagenomic sequences as input, scans through each input DNA sequence, and counts the number of co-occurrences of adjacent *k*-mer pairs <*k*-meri ><*k*-mer_2_> using a sliding window of length *2k*. Given the alphabet Σ = {*A, C, G, T*}, the resultant *k*-mer co-occurrence matrix can be recorded as *X* ∈ ℝ^|Σ|^*k*^ × |Σ|^*k*^^, where *X_ij_* indicates the count of *k*-mer *i* observed in precedence to *k*-mer *j*.

A desirable *embedding* model should possess the following two properties: (i) preserving the co-occurrences between pairwise *k*-mers; (ii) since each *k*-mer has its own bias originated from sequence background noise, it is also desirable to decouple the observed *k*-mer counts into sequence-specific background noise and its true signals. Mathematically, for each *k*-mer *i*, the objective is to find the biases 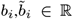 and *d*-dimensional vectors 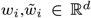 for each *k*-mer *i*, where *w_i_* and 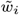 correspond to the vectors of *k*-mer *i* appearing in the preceding and succeeding position of the co-occurrence pairs, respectively. The cooccurrence count *X_ij_* between *k*-mer *i* and *k*-mer *j* can be approximated by:

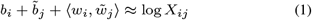

where *X_ij_* is log-transformed to avoid the skewed distribution of nonnegative co-occurrences (Landauer, 2006). *b_i_* and 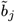 are sequencespecific biases, representing the background noise estimated from the sequence per se. The motivation of Equation 1 is to decouple the observed *k*-mer count into the background noise and the true signal (Lu *et al.*, 2017a; Ren *et al.*, 2018), which is similar to 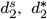, and CVTree. And 〈±,±〉 indicates the inner product between two vectors, that is, 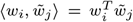. Note that the asymmetry holds between *X_ij_* and *X_ji_* because 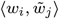 and 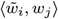 are distinct. Instead of parameterizing the very high-dimensional *k*-mer co-occurrence matrices, the *embedding* model performs a low-rank approximation by factorization. Equation 1 is optimized by least squares regression in the following form:

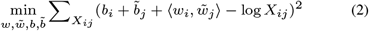

The representation obtained by Equation 2 for *k*-mer *i* reduces the *i*-th row size in the co-occurrence matrix from *n* = 4^*k*^ to 2d + 2. In the setting when *k* = 4 and *d* =10, the human genome is compressed to 52 KB from 3.1 GB.

### 2.3 CRAFT *dissimilarity*

Let *X_i_*. = Σ_*j*_ *X_ij_* be the occurrence of *k*-mer *i* preceding any *k*-mer pairs. Additionally, let *P_ij_* = *X_ij_ / X_i_*. be the conditional probability that *k*-mer *j* is observed right after the observation of *k*-mer *i*. *P_ij_* can be represented by 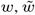 and *b* obtained by Equation 2:

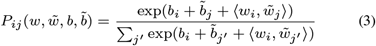

It is worth mentioning that Equation 3 can be interpreted as performing a softmax operation across all succeeding *k*-mers.

A desirable *dissimilarity* model aims to quantitatively compare two genome sequences *G*^(1)^ and *G*^(2)^, based upon their *embedding* models 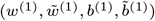 and 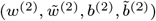, respectively. Let 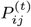 be the conditional probability defined in Equation 3, where *t* = 1, 2. As suggested by Lu *et al.* (2017a), the dissimilarity between *G*^(1)^ and *G*^(2)^ is defined as:

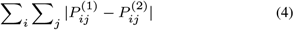

### 2.4 Acceleration in searching massive repositories

In the case when the archived repository is huge, exhaustive searching every genome in the repository is inevitably laborious. Therefore, CRAFT uses a branch-and-bound strategy to accelerate the searching. Specifically, CRAFT first identifies top-*K genus* in the sequence repository *D*, and sequently search candidate species from these *genus* only. Say for each *genus g*, let *D_g_* ⊂ *D* be the sequences sharing the same *genus g*, CRAFT constructs a smaller representative set of *g*, 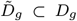, by randomly choosing 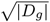 sequences from *D_g_*. In doing so, identifying top-*K genus* reduces searching from *D* into 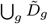. For example, in the NCBI Assembly repository with 139, 576 genome sequences, the size of 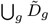 is 10,484 and the size of overall searched species is 7,419, respectively. The software architecture was given in Supplementary Note1.

## 3 Results

### 3.1 The compression ratio of common repositories

CRAFT is used to archive three widely-used genome/metagenome repositories, including NCBI Assembly with 139, 576 genome sequences, RefSeq representative genomes with 7,106 genome sequences, and the Human Microbiome Project 1-II (HMP 1-II) with 2, 355 metagenome samples with *d* =10 and *k* = 4. See Table 1 and Supplementary Note 2 for detailed summery of repositories. By empirically using *k* = 4, CRAFT reduces the size of three massive repositories from 829.60GB (.fa), 376.38GB (.fa), and 7.13TB (.sra) to 2.93GB, 152.67MB, and 50.40MB, respectively.

**Table 1.** The summary of three genome/metagenome repositories

| Repository | Genome number | Repository size | Compressed size | Compression ratio |
| --- | --- | --- | --- | --- |
| NCBI Assembly | 139,576 | 829.60GB(.fa) | 2.93GB | 3.5‰ |
| RefSeq representative | 7,106 | 376.38GB(.fa) | 152.67MB | 0.4‰ |
| HMP 1-II | 2,355 | 7.13TB(.sra) | 50.40MB | 0.0067‰ |

### 3.2 The dissimilarity estimations between genome/metagenome sequences

We evaluated CRAFT in dissimilarity estimation among genomes or metagenomes on real datasets: 28 vertebrate genomes, 21 primate genomes, 27 *E.coli/Shigella* genomes, and 28 mammal gut metagenome samples. As shown in Figure 2, the genomes among 28 vertebrate genomes or 28 mammal gut metagenomic samples are more dissimilar to each other than those among 27 *E.coli/shigella* genomes and 21 primate genomes, which is consistent across three different measures, 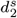, Mash, and Skmer. CRAFT is compared to six state-of-the-art alignment-free dissimilarity measures including four *k*-mer frequency-based methods: Manhattan, CVTree, 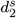, and 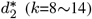; and two *k*-mer hashing-based methods: Mash and Skmer (*k*=12~31). For 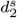 and 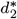, the Markov order is selected based on BIC (Narlikar *et al.*, 2012). For Mash and Skmer, the default sketch size is applied, that is, 10^7^ for Skmer and 10^3^ for Mash. The embedding dimensionality *d* is set to 10 throughout the experiments, and see Supplementary Note 1 for the insensitivity of choosing *d*. The performance is investigated by the correlations between estimated dissimilarity and evolutionary distance, or the distance between the reference and the clustered tree.

**Fig. 2.**
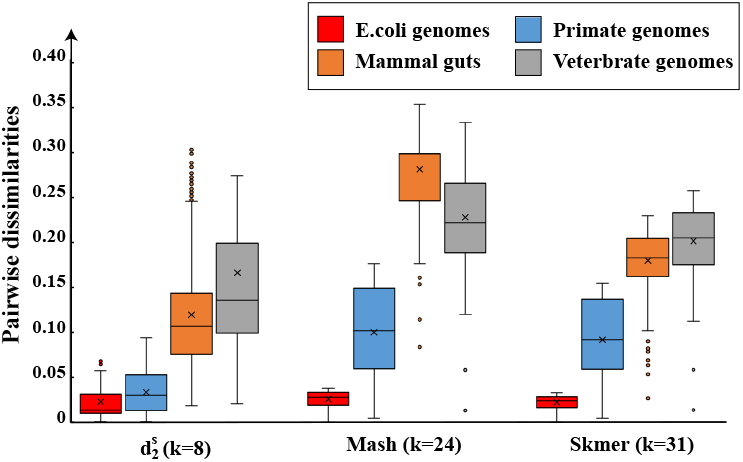
The pairwise dissimilarities of genomes among four datasets, calculated by 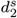, Mash, and Skmer.

#### 3.2.1 The vertebrate and the primate genome dataset

We used CRAFT to estimate the pairwise dissimilarities among 28 vertebrate genomes, as well as 21 primate genomes (Miller *et al.*, 2007). The estimated dissimilarities are compared against the corresponding evolutionary distances calculated by *Ape* (Paradis *et al.*, 2004), in terms of Spearman correlations. As shown in Figure 3, in the 28 vertebrate genome dataset of diverse dissimilarity, CRAFT outperforms other state-of-the-art measures. In particular, CRAFT performs comparably to Mash by using much smaller *k*-mer size (*k*=8 versus *k*=31). However, in the 21 primate genome dataset of high similarity (Figure 2), all *k*-mer frequency-based methods, including CRAFT, are not as sensitive as hashing-based methods such as Mash and Skmer, which use much longer *k*-mer size.

**Fig. 3.**
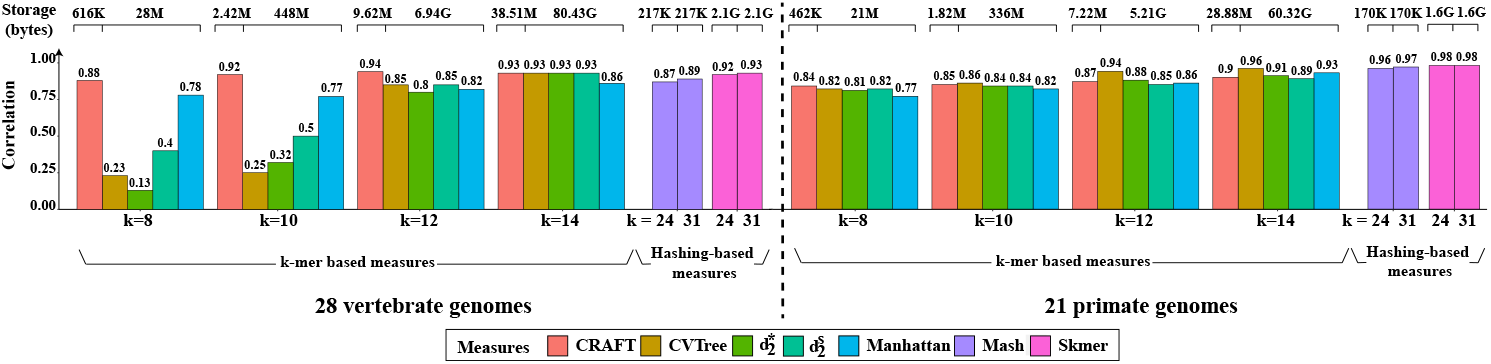
The Spearman correlation between the pairwise evolutionary distances and the dissimilarities among the 28 vertebrate genomes and 21 primate genomes calculated by CRAFT and various dissimilarity measures. The storage of each method is presented at the top.

Additionally, CRAFT has notable advantages over competing methods in running time, particularly for short *k*-mers, even comparable to hashing-based methods such as Skmer and Mash, as illustrated in Figure 4. It is worth mentioning that CRAFT achieves such performance with only 38.51MB storage consumption, in contrast to 80.43GB 14-mer counts for CVTree, 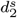 and 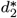. See Figure 3 for detailed comparison of storage consumption. In short, CRAFT consistently outperforms state-of-the-art *k*-mer frequency-based methods, with much succinct storage usage. When compared to state-of-the-art hashing-based methods such as Skmer and Mash, the superiority of CRAFT is not guaranteed and dependent on the data analysis task.

**Fig. 4.**
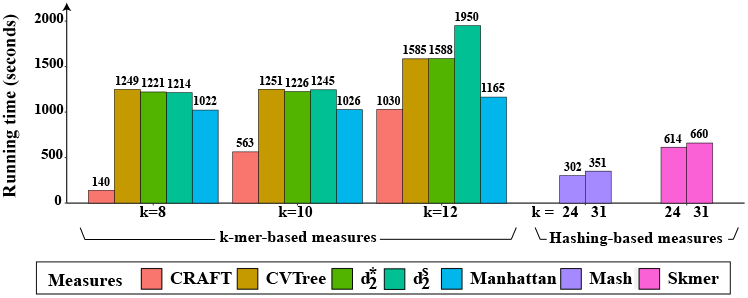
Running time (seconds) used in calculating the dissimilarities among 28 vertebrate genomes.

#### 3.2.2 The mammal gut metagenome samples dataset

We used CRAFT to analyze a mammalian gut metagenome dataset consisting of 28 HTS samples (Muegge *et al.*, 2011). These samples split into 3 groups, including 8 hindgut-fermenting herbivores, 13 foregut-fermenting herbivores and 7 simple-gut carnivores. We investigated how well CRAFT can identify these groups in comparison against other alignment-free dissimilarity measures.The experimental settings follows Section 3.2.1 for sketch size and the Markov order. For every measure,we used UPGMA (Murtagh, 1984) to cluster the 28 samples based upon the calculated dissimilarity matrix, with each group colored differently.

As shown in Figure 5(A), CRAFT achieves the best separations among 3 groups, compared to the other six measures. The superior performance of CRAFT is quantitatively supported by the symmetric difference shown in Figure 6. The symmetric difference (Robinson and Foulds, 1981) is the quantitative metric to evaluate the consistency between two trees, where the lower symmetric difference indicates the better consistency, calculated *Treedist* from *Phylip* (http://evolution.genetics.washington.edu/phylip.html). It is worth mentioning that even within the same group, CRAFT tends to cluster the same species together, such as the lions in the group of simplegut carnivores, which is failed by CVTree and 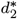. In short, with much less storage consumption, as depicted in Figure 6, CRAFT is the best performer. The hierarchical clustering trees obtained by other 30 alignment-free measures were given in Figure S3 of Supplementary Note 1. Analogous to the veberate genome dataset, CRAFT shows again the advantages in distinguishing genomes of diverse dissimilarities.

**Fig. 5.**
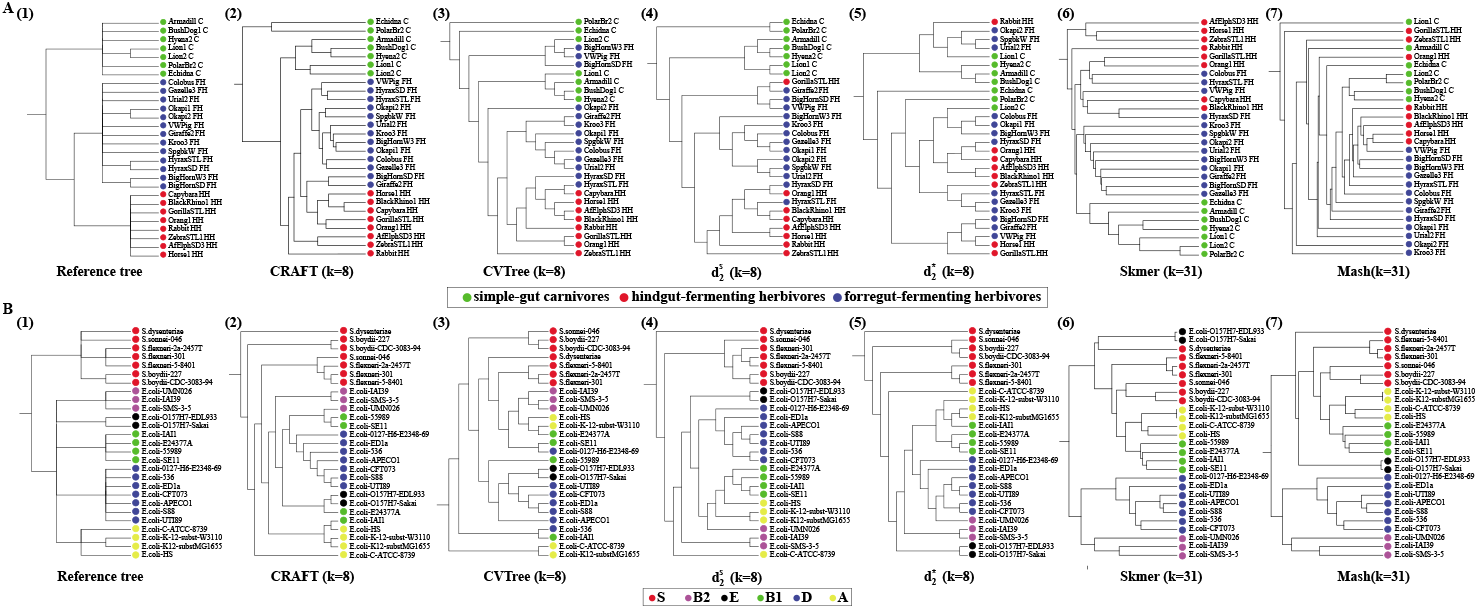
The clustering results of (A) 27 *E. coli* and *Shigella* genomes, and (B) 28 metagenome samples from mammal gut using CRAFT, CVTree, 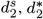, Mash, and Skmer.

**Fig. 6.**
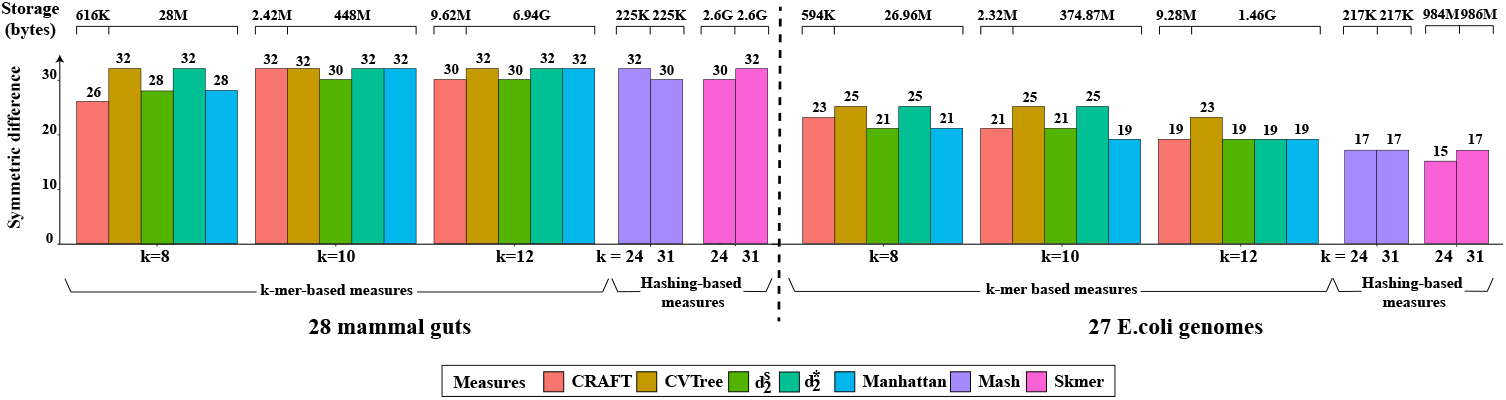
Symmetric difference between the clustering and reference trees on mammal gut metagenome dataset and *E.coli* genome dataset. The lower of symmetric difference is, the more similar the clustering and reference trees are. The storage of each method is presented at the top.

#### 3.2.3 The *E. coli* and *Shigella* genomes dataset

We also applied CRAFT to analyze the genomic sequences of 20 *E. coli* and 7 *Shigella* genomes (Bernard *et al.*, 2016). These genomes are categorized into six *E. coli* reference (ECOR) groups: A, B1, B2, D, E, and S.

As shown in Figure 5(B), for CRAFT, each ECOR is monophyletic except B1 and E with respect to the reference tree. As shown in Figure 6, analogous to the 21 primate genome dataset, all *k*-mer frequency-based methods, including CRAFT, are not as sensitive as hashing-based methods such as Mash and Skmer, which use much longer *k*-mer size. However, CRAFT is still the best performer among all *k*-mer frequency-based methods. See Figure S2 in Supplementary Note 1 for the hierarchical clustering trees obtained by other 30 alignment-free measures.

### 3.3 Query HTS genomic data in the compressed genomic sequence databases

In this section, CRAFT is used to search random HTS genomic and metagenomic samples against archived massive sequence repositories, including NCBI Assembly, Refseq representative and HMP 1-II. To evaluate the performance of CRAFT thoroughly, the testing HTS samples are intentionally chosen from different major taxonomic branches. Note that mitochondrial sequences are excluded to search against the NCBI Assembly database and the Representative Genomic database based upon the fact that mitochondrial sequences have different *k*-mer frequencies from chromosomal sequences (Mrázek and Karlin, 2007). Similarly, ribosome RNA sequencing data is also excluded to search against HMP 1-II.

#### 3.3.1 Query HTS genomes in the Refseq Representative database

The HTS samples corresponding to 104 random species are queried against the RefSeq representative repository. The selected species are intentionally chosen to cover 8 major taxonomic branches, including archaea, bacteria, fungi, invertebrate, plant, protozoa, vertebrate mammalian, and vertebrate other. As shown in Figure 7, CRAFT is able to pinpoint 90 out of 104 queried HTS samples to their true species within top 2 matches. We use a state-of-the-art alignment-free measure, Manhattan distance, to query the remaining 14 failed-matched HTS samples, to investigate whether these failures are exclusive to CRAFT. As shown in Supplementary Note 3, Manhattan distance achieved comparable performance with negligible difference, suggesting that CRAFT suffers the same limitations as general alignment-free measures. Without nucleotide by nucleotide alignments, it is reasonable that alignment-free methods cannot achieve full accuracy as alignment-based methods.

**Fig. 7.**
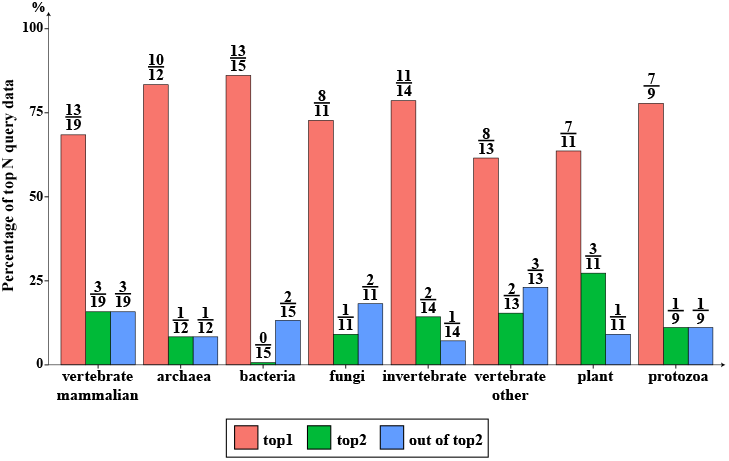
The performance of CRAFT in the query of 104 HTS data in Refseq representative database. The 104 HTS data from 8 major taxonomies were tested in CRAFT. For each major taxonomy, the distribution of the source genome’s ranking in the matching list were given. Totally, 77 data out of the 104 are matched to their source genome as the top one in the matching list.

#### 3.3.2 Query E. *Coli* strains in the NCBI Assembly database

The HTS samples corresponding to three random *E. Coli* strains are queried against NCBI Assembly repository, including *E. Coli* O157:H7, *E. Coli* O26:H11, and *E. Coli* O111. As shown in Table 2, CRAFT is able to distinguish the subtle differences among various strains from the same *E. Coli* species. Specifically, two out of three *E. Coli* strains, *E. Coli* O157:H7 and *E. Coli* O26:H11 are matched exactly to their source genomes and the remaining one, *E. Coli* O111 can also be matched its source genome within top 4 matches. Additionally, it is worth mentioning that all top 10 matches of the three strains belong to the targeted *E. Coli* species, showing the great sensitivity of CRAFT.

**Table 2.** Queries of HTS data from three *E. Coli* strains by CRAFT

| Organism | *E.coli* O157:H7 (SRR6297305) |  | *E.coli* O111 (SRR6164651) |  | *E.coli* O26:H11 (SRR6373714) |  |
| --- | --- | --- | --- | --- | --- | --- |
| Ranking | Species | Dissimilarity | Species | Dissimilarity | Species | Dissimilarity |
| 1 | *E.coli* O157:H7 strain=EC4486 | 18.3836 | *E.coli* strain=211 | 20.1988 | *E.coli* O26:H11 strain=11368 | 19.4856 |
| 2 | *E.coli* O26:H11 strain=12KH23 | 18.4500 | *E.coli* strain=GZC12-13 | 20.2132 | *E.coli* strain=12-269M | 19.5676 |
| 3 | *E.coli* strain=22593 | 18.5616 | *E.coli* strain=13f2-1 | 20.5486 | *E.coli* O26 strain=RM10386 | 19.6440 |
| 4 | *E.coli* strain=211 | 18.6236 | *E.coli* O111:H- strain=11128 | 20.5607 | *E.coli* strain=Ecol_AZ146 | 19.7212 |
| 5 | *E.coli* strain=13f2-1 | 18.6419 | *E.coli* strain=11a5 | 20.5843 | *E.coli* O26 strain=RM8426 | 19.8673 |
| 6 | *E.coli* strain=11a2 | 18.7134 | *E.coli* O26:H11 strain=12KH23 | 20.5894 | *E.coli* strain=MDR_56 | 19.8766 |
| 7 | *E.coli* strain=11a5 | 18.7244 | *E.coli* strain=MOD1-EC2815 | 20.6058 | *E.coli* strain=2013C-3277 | 19.9180 |
| 8 | *E.coli* strain=FDAARGOS_398 | 18.9112 | *E.coli* | 20.6529 | *E.coli* strain=2013C-4225 | 19.9578 |
| 9 | *E.coli* strain=13h3 | 18.9969 | *E.coli* 26:H11 strain=PV1125 | 20.6659 | *E.coli* strain=UMNK88 | 20.0086 |
| 10 | *E.coli* strain=EXF1-6_2013 | 19.0427 | *E.coli* strain=11a2 | 20.6949 | *E.coli* strain=NCTC8783 | 20.0757 |

#### 3.3.3 Query of HTS metagenomes data in HMP 1-II

The HTS samples corresponding to 13 random metagenomes are queried against the HMP 1-II repository, covering different parts of the human body such as skin, vagina, gut and oral cavity. As shown in Supplementary Note 3, 11 out of 13 samples can be exactly matched to their targeted body locations. For example, the metagenome from the human skin of retroauricular crease (SRR3182045) is not only matched to the correct source of microbial community (i.e., human skin), but pinpoint to its exact retroauricular crease location as well.

### 3.4 Robustness for incomplete data

In this section, we investigate the robustness of CRAFT by exploring its limit in extreme cases. Specifically, we demonstrate that CRAFT can still obtain accurate searching performance even when the query genome sequences are of low coverage or the HTS data is of very low sequence depth.

#### 3.4.1 Low coverage of genome sequence

For each of the eight major taxonomic branches, we randomly selected five genomes. Each genome was randomly cut out a fragment/fragments which cover the 80%-1% of the whole genome. And the fragement(s) were set as a query data to see whether CRAFT can find the source genome under the specified coverage proportion of the whole genome. The Figure 5 shows the numbers which can find the source genome at the best match under different coverage of the whole genome. The experiment shows that the proportion of the sequence to the complete genome has notable effect on the matching performance. As shown in Figure 8 and Supplementary Note 3, all the fragments covering more than half of the whole genome can find the exact source genome at the best match in most case. While, even for the 2% coverage for the whole genome sequences, there are still more than half of the testing data can find their source genomes. It is reasonable because longer sequence has more consistent distribution with the complete genome than the short sequence.

**Fig. 8.**
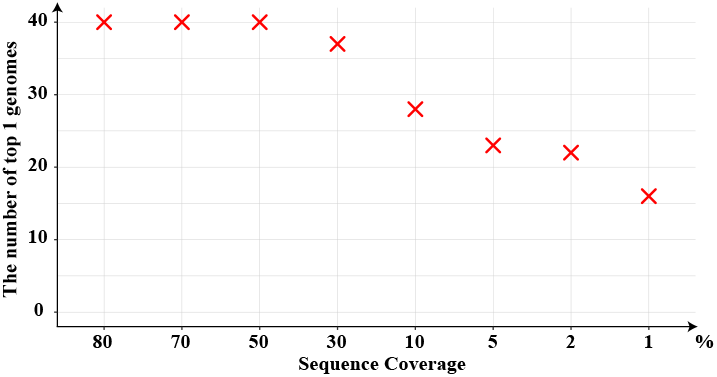
The performance of CRAFT for incomplete genome data. The sequence was randomly selected for 40 genomes from the eight major taxonomic branches and set as the query data to CRAFT. The horizontal axis is the proportion of the selected sequence to the whole genome. the vertical axis is the number of data which can find their source genome at the best match under different sampling rates among the 40 genomes.

#### 3.4.2 Low sequence depth of HTS data

For each of the eight major taxonomic branches, five HTS data was randomly selected. For each HTS data, the reads were randomly sampled with the rate from 80% to 0.001% as a query data with different sequencing depth to test whether CRAFT can find the source genome at the best match. In Figure 9, the vertical axis is the number of data which can find their source genome at the best match under different sampling rates among the 40 HTS data. Even keeping only 5% of reads, CRAFT still can embed the sequence into the vector space accurately and hit the source genome at the best match for all of the 40 testing data. Being different with fragments in complete genome, random sampled reads still cover the genome region evenly and therefore reflect the *k*-mer distribution accurately.

**Fig. 9.**
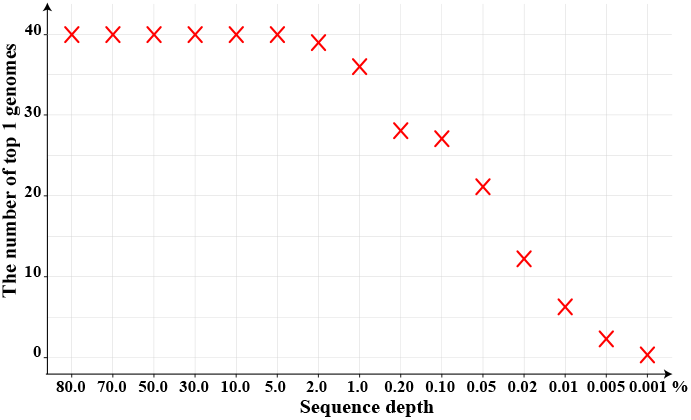
The performance of CRAFT for sampled HTS data. Forty HTS data from the eight major taxonimc branches were sampled with rate from 80% to 0.001%. The number of sampled data which can find their source genome at the best match decreases as the sampling rate decreases. And for all of the 40 testing data, even only keeping 5% reads, CRAFT can still find the source genome at the best match.

## 4 Conclusion and discussions

In this study, we developed CRAFT, a stand-alone tool for compact representations of gigantic sequence databases and fast query of genomic/metagenomic sequence, with user-friendly graphical interface with one-click installation. From the practitioner’s perspective, CRAFT not only integrates three built-in common repositories, but support three types of downstream visualized analysis as well. In summary, with compact storage of massive sequence repositories, CRAFT preserves the sequence comparison accuracy compared to other state-of-the-art alignment-free measures such as Manhattan distance, CVTree, 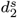, and 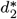. We are also aware that CRAFT suffers the same limitation of *k*-mer frequency-based methods, that is, the usage of relative short *k*-mers is not as powerful as long *k*-mers in capturing the subtle difference among extremely similar sequences. Extending CRAFT to cope with much longer *k*-mers would be an interesting direction for future research. CRAFT is designed for extensibility. Though CRAFT has only archived repositories containing genomic/metagenomic DNA sequences so far, in principle it would be applicable to other types of data, ranging from genomes, metagenomes, transcriptomes and metatranscritpomes, as long as DNA sequences, RNA sequences, or amino acid sequences can be represented as *k*-mers

