## Supplemental Note 1 for "CRAFT: Compact genome Representation towards large-scale Alignment-Free daTabase"

**Supplementary Note 1**

1. **Software architecture of CRAFT**

The software architecture of CRAFT is composed of backend and frontend. As shown in Figure S1, the backend of CRAFT composes of four modules and written with c++, where *kmc* and *glove* are the existing tools for *k-mer* counting and embedding. The frontend designs a visual user interface and displays the results on different operating system. The frontend is written with HTML, CSS and JavaScript, and runs on *node.js* and *Electron*.

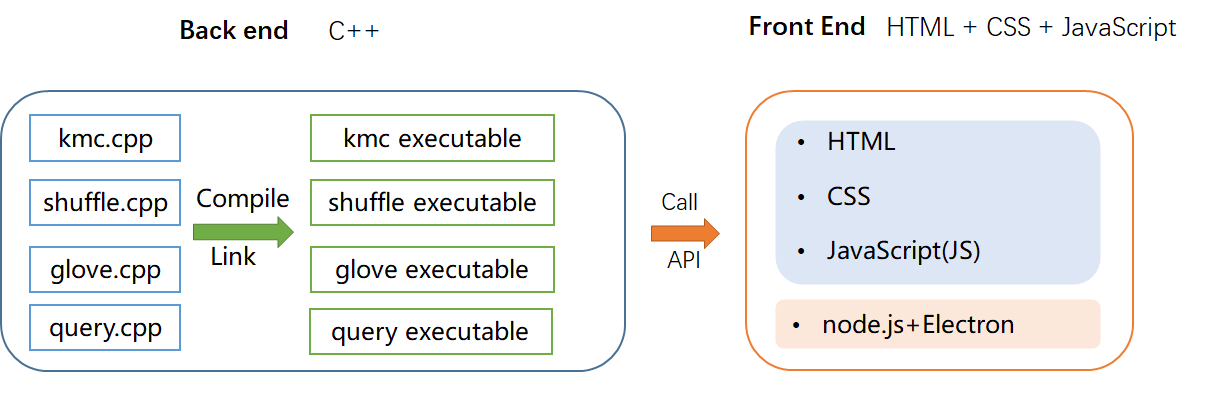

Figure S1 The software architecture of CRAFT

2. **CRAFT and 30 alignment-free measures on 27 E.coli/Shigella microbial genomes and 28 mammal guts metagenomic samples**

We compared the dissimilarities calculated by CRAFT and 28 other alignment-free measures calculated by CAFE [1] on 27 *E. coli* and *Shigella* genomes [2], which are categorized into six *E. coli* reference (ECOR) groups. The hierarchical clustering tree was built with UPGMA [3] on the calculated pairwise dissimilarity matrix. The $k$-mer length was set as 8 for k-mers based tool and 31 for Mash and Skmer. The clustering trees are shown in Figure S2.

CRAFT was also applied to analyze a mammalian gut metagenomic dataset [4] comprised of high throughput sequencing reads from 28 metagenomic samples (i.e., 8 hindgut-fermenting herbivores, 13 foregut-fermenting herbivores and 7 simple-gut carnivores). Identical experimental setting was applied. The clustering results from CRAFT and 28 other alignment-free measures are given in Figure S3.

3. **The selection of parameter *d* in the performance of CRAFT on the four datasets**

The embedding dimension *d* in our model is generally set as 5~15. Our experiments in different values of *d* show that *d* has no much effect on the results, especially with the increase of length of *k*.

Table S1. Spearman Correlation for ANI and CRAFT dissimilarity with different values of *d* for 28 vertebrates

| **CRAFT** | **d=5** | **d**=**10** | **d**=**15** |
| --- | --- | --- | --- |
| **k=8** | 0.88 | 0.85 | 0.84 |
| **k=10** | 0.92 | 0.92 | 0.91 |
| **k=12** | 0.93 | 0.93 | 0.94 |

Table S2. Spearman Correlation for ANI and CRAFT dissimilarity with different values of *d* for 21 primates

| **CRAFT** | **d=5** | **d**=**10** | **d**=**15** |
| --- | --- | --- | --- |
| **k=8** | 0.84 | 0.84 | 0.84 |
| **k=10** | 0.84 | 0.85 | 0.85 |
| **k=12** | 0.86 | 0.87 | 0.87 |

Table S3. Symmetric difference for reference and hierarchical clustering tree of CRAFT with different values of *d* for 28 mammal guts

| **CRAFT** | **d=5** | **d**=**10** | **d**=**15** |
| --- | --- | --- | --- |
| **k=8** | 26 | 28 | 28 |
| **k=10** | 32 | 32 | 32 |
| **k=12** | 30 | 30 | 30 |

Table S4. Symmetric difference for reference and hierarchical clustering tree of CRAFT with different values of *d* for 27 E.coli/Shigella genomes

| **CRAFT** | **d=5** | **d=10** | **d=15** |
| --- | --- | --- | --- |
| **k=8** | 27 | 23 | 23 |
| **k=10** | 21 | 21 | 21 |
| **k=12** | 21 | 19 | 19 |

4. **The Robinson-Foulds (RF) score and Symmetric difference between reference and hierarchical clustering tree of CRAFT on the 28 mammal guts and 27 E.coli/Shigella genomes**

Besides Symmetric difference, Robinson-Foulds is also calculated, which is another measure for the dissimilarity between two tree topologies with the same number of leaves and the same labels (species) at the leaves, i.e., it measures the dissimilarity of branching patterns and ignores branch lengths. The results are shown in Table S5,S6 for the two datasets. For easy comparison, we also give symmetric difference. It is clear that the two measures demonstrate similar evaluation results. And because Robinson-Foulds ignores branch lengths, it will miss some information in some cases.

**Table S5. Robinson-Foulds and Symmetric difference for reference and hierarchical clustering tree of CRAFT and other measures for 28 mammal guts**

| **k** | **8** | **10** | **12** | **14** | **21** | **24** | **31** |
| --- | --- | --- | --- | --- | --- | --- | --- |
| **Robinson-Foulds** | | | | | | | |
| **CRAFT** | 28 | 34 | 32 | NA | NA | NA | NA |
| **Mash** | NA | NA | 36 | 34 | 34 | 34 | 34 |
| **Skmer** | NA | NA | 36 | 34 | 34 | 34 | 32 |
| **cvtree** | 34 | 34 | 34 | NA | NA | NA | NA |
| $\boldsymbol{d}_{\boldsymbol{2}}^{\boldsymbol{s}}$ | 28 | 32 | 32 | NA | NA | NA | NA |
| $\boldsymbol{d}_{\boldsymbol{2}}^{\boldsymbol{*}}$ | 32 | 34 | 34 | NA | NA | NA | NA |
| **Mahatten** | 30 | 34 | 34 | NA | NA | NA | NA |
| **Symmetric Difference** | | | | | | | |
| **CRAFT** | 26 | 32 | 30 | NA | NA | NA | NA |
| **Mash** | NA | NA | 32 | 32 | 32 | 32 | 30 |
| **Skmer** | NA | NA | 32 | 32 | 32 | 30 | 32 |
| **cvtree** | 32 | 32 | 32 | NA | NA | NA | NA |
| $\boldsymbol{d}_{\boldsymbol{2}}^{\boldsymbol{s}}$ | 26 | 30 | 30 | NA | NA | NA | NA |
| $\boldsymbol{d}_{\boldsymbol{2}}^{\boldsymbol{*}}$ | 30 | 32 | 32 | NA | NA | NA | NA |
| **Mahatten** | 28 | 32 | 32 | NA | NA | NA | NA |

**Table S6. Robinson-Foulds and Symmetric difference for reference and hierarchical clustering tree of CRAFT and other measures for 27 E.coli/Shigella genomes**

| **k** | **8** | **10** | **12** | **14** | **21** | **24** | **31** |
| --- | --- | --- | --- | --- | --- | --- | --- |
| **CRAFT** | 25 | 21 | 21 | NA | NA | NA | NA |
| **Mash** | NA | NA | 21 | 19 | 19 | 19 | 19 |
| **Skmer** | NA | NA | 21 | 19 | 19 | 19 | 19 |
| **cvtree** | 27 | 27 | 25 | NA | NA | NA | NA |
| $\boldsymbol{d}_{\boldsymbol{2}}^{\boldsymbol{s}}$ | 23 | 23 | 21 | NA | NA | NA | NA |
| $\boldsymbol{d}_{\boldsymbol{2}}^{\boldsymbol{*}}$ | 27 | 27 | 21 | NA | NA | NA | NA |
| **Mahatten** | 23 | 21 | 21 | NA | NA | NA | NA |
| **Symmetric Difference** | | | | | | | |
| **CRAFT** | 23 | 21 | 19 | NA | NA | NA | NA |
| **Mash** | NA | NA | 19 | 17 | 17 | 17 | 17 |
| **Skmer** | NA | NA | 19 | 17 | 17 | 15 | 17 |
| **cvtree** | 25 | 25 | 23 | NA | NA | NA | NA |
| $\boldsymbol{d}_{\boldsymbol{2}}^{\boldsymbol{s}}$ | 21 | 21 | 19 | NA | NA | NA | NA |
| $\boldsymbol{d}_{\boldsymbol{2}}^{\boldsymbol{*}}$ | 25 | 25 | 19 | NA | NA | NA | NA |
| **Mahatten** | 21 | 19 | 19 | NA | NA | NA | NA |

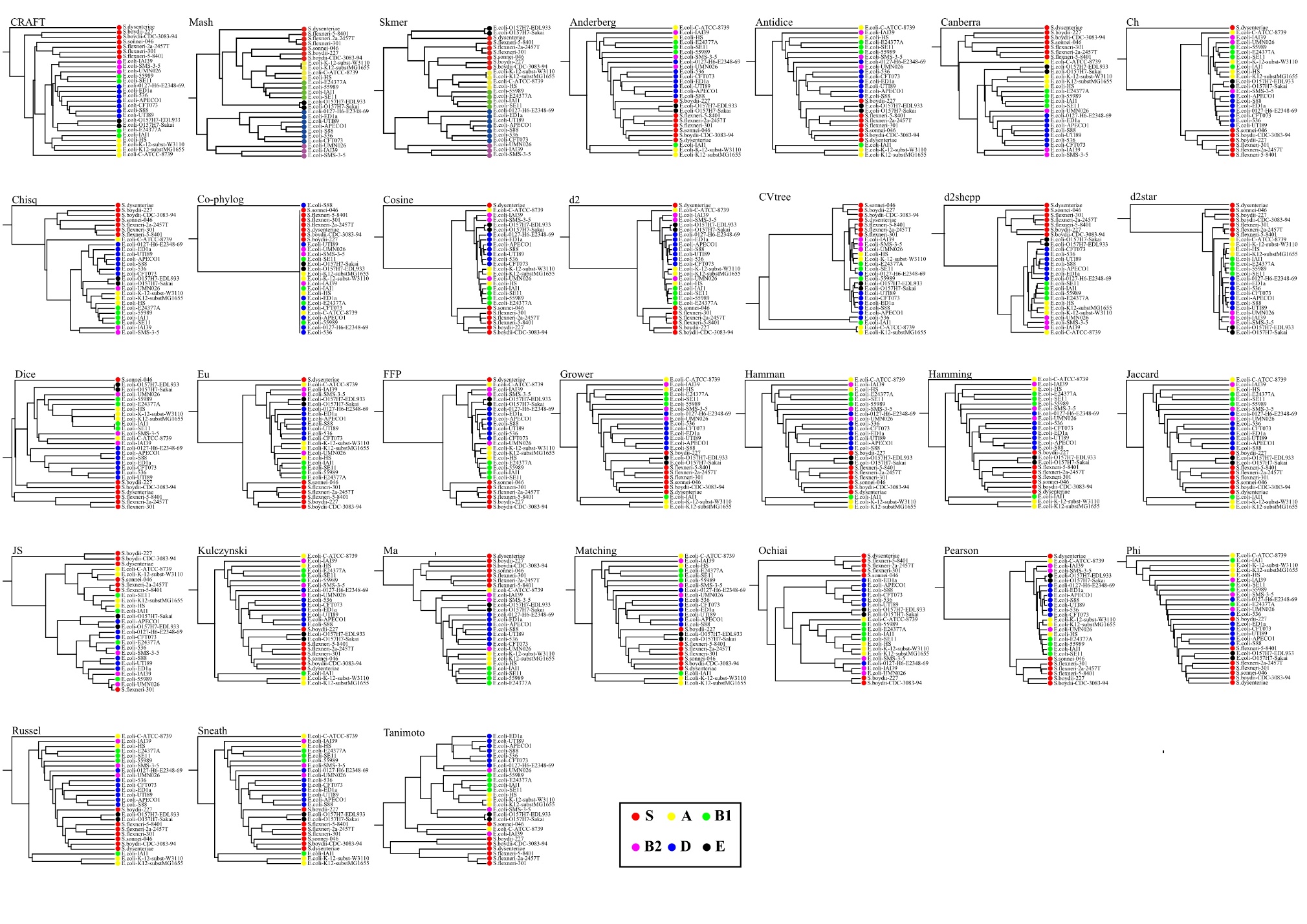

Figure S2 Clustering tree using CRAFT and 28 other alignment-free measures on 27 E.coli/Shigella genomes

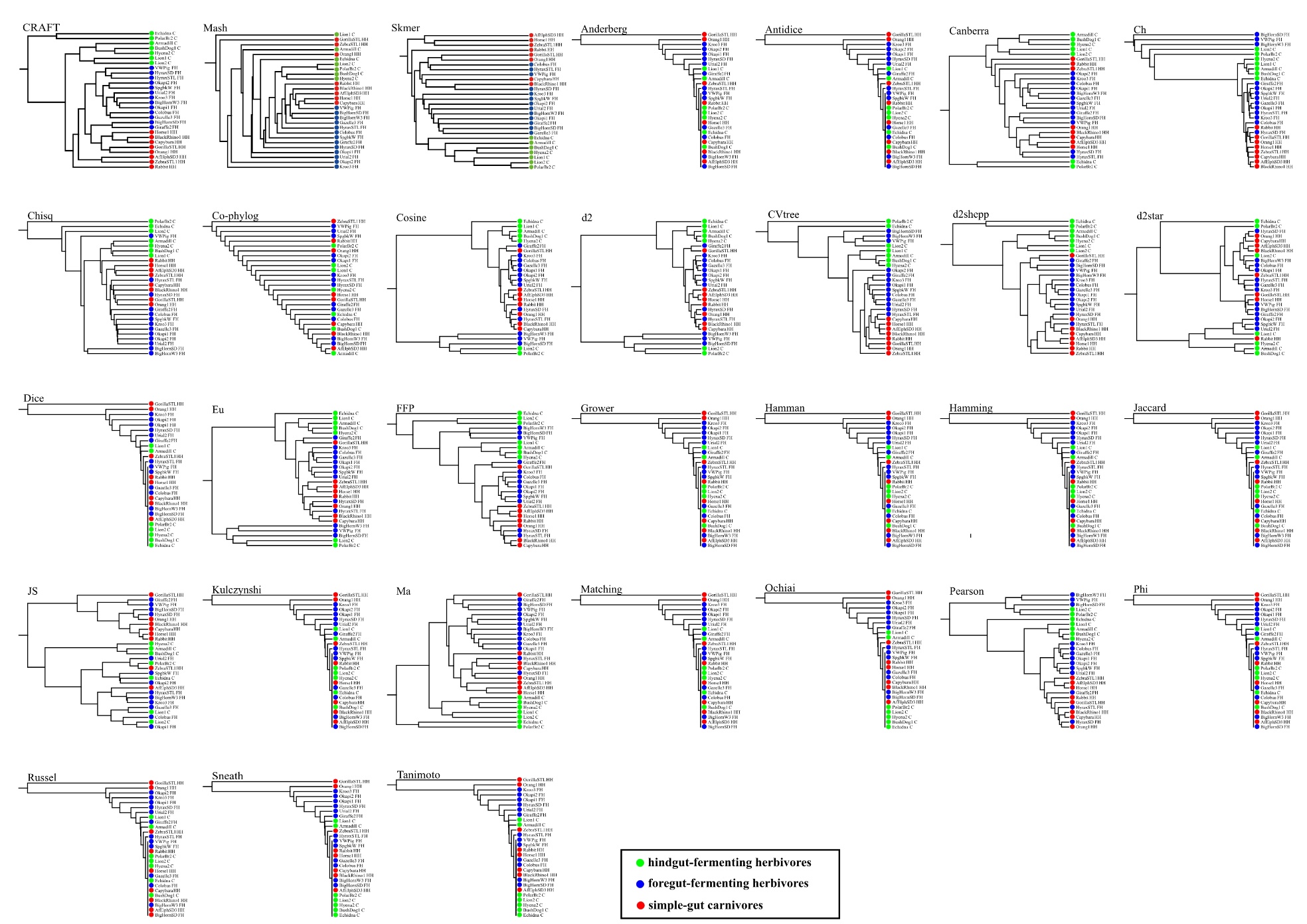

Figure S3 Clustering tree using CRAFT and 28 other alignment-free measures on 28 metagenomic samples
